# Experimental speciation: selection against hybridization strengthens reproductive barriers and limits introgression

**DOI:** 10.64898/2026.10.15.757478

**Authors:** Graham C. McLaughlin, Verónica Baquero-Méndez, Sophia Sudak, Brian Hollis

## Abstract

Evolutionary theory predicts that when hybridization between divergent populations is costly, natural selection should favor traits that reduce interbreeding, a process known as reinforcement^1–3^. Direct experimental evidence of reinforcement is rare, however, and no study has demonstrated that reinforcing selection can strengthen reproductive isolation between naïve populations while allowing continuous gene flow^4,5^. Here we show that >100 generations of experimental selection against hybridization – imposed by removing a controlled fraction of GFP-marked hybrids – led to the evolution of increased reproductive isolation between two populations of *Drosophila melanogaster* brought into secondary contact. Selection strengthened both sexual isolation (via reduced female remating with migrant males) and postmating-prezygotic isolation, although the specific barriers that evolved depended on the strength of selection against hybridization. Sexual isolation was evident only when mated females encountered both within-population and migrant males, revealing an important role for social mating context. Notably, intermediate selection against hybridization produced the strongest overall increase in reproductive isolation, suggesting that reinforcement is most effective when maladaptive hybridization with both newly arrived immigrants and early-generation hybrids remains sufficiently common to generate persistent selection for avoidance. Whole-genome sequencing revealed that divergence was concentrated in regions linked to the GFP barrier locus, where introgression was reduced. These regions were also enriched for loci showing signatures consistent with an adaptive response to selection against hybridization. Together, these results provide direct experimental evidence that reinforcement can increase reproductive isolation under ongoing gene flow and reveal how selection against hybridization shapes genomic divergence during the early stages of speciation.

## Main text

Speciation requires the evolution of reproductive isolation (RI) and its persistence in the face of gene flow. When diverging populations come into secondary contact, hybridization is expected to erode reproductive barriers, potentially leading to fusion rather than speciation^6–8^. Yet, reproductive barriers are often maintained and sometimes strengthened despite ongoing gene flow^9–12^, implying that selection can actively counter homogenization. One mechanism that could produce this outcome is reinforcement, in which selection against maladaptive hybridization favors traits that reduce interbreeding between populations^1–3,13–15^. If hybrid matings produce fewer or less fit offspring relative to conspecific matings, individuals that avoid hybridization gain a fitness advantage, leading to the evolution of increased assortative mating and stronger reproductive isolation^13,16^.

Despite a long theoretical history^13,17–19^ and its conceptual appeal, direct demonstrations of reinforcement remain rare^5^. Comparative approaches have provided suggestive evidence, most notably the pattern of reproductive character displacement (RCD)^20^, in which prezygotic reproductive isolation is stronger between taxa in sympatry than in allopatry^21–30^. Although this geographic pattern is consistent with reinforcement, it can also arise through ecological divergence or differential fusion and extinction^31–33^, making it difficult to distinguish among these mechanisms^26^. Only a few systems provide strong support for reinforcement while ruling out alternative explanations^34–38^. Other studies have failed to confirm reinforcement as the driver of increased RI in cases where it was initially suspected^16,39–41^. Therefore, inferring reinforcement from comparative data alone is challenging, and the causal role of selection against hybridization often cannot be established after the fact^13,15,31,42^.

Experimental evolution, which involves manipulating replicated populations and tracking evolution in real time^43^, provides a powerful approach to directly test whether reinforcing selection can strengthen reproductive isolation. In this context, secondary contact can be recreated in the laboratory while the fitness consequences of hybridization are experimentally controlled, for example by removing hybrid offspring each generation^4–6,44,45^. Experimental tests of reinforcement between naïve populations are rare, and none have successfully strengthened reproductive isolation while allowing continuous gene flow^4,45^. Past experiments removed all hybrids, thereby preventing gene flow entirely^4,6^, or involved distinct species with prior evolutionary histories of reinforcing selection^36,46,47^. Empirical studies investigating the genomic architecture of reinforcement are also limited^34,48,49^ and no experimental reinforcement study has examined genome-wide responses to selection^5,44^. Here, we leverage the power of experimental evolution to directly test whether selection against hybridization can strengthen reproductive isolation under ongoing gene flow and characterize how reinforcing selection shapes the genome.

### A laboratory system for experimental reinforcement

The IV and LH_M_ laboratory lineages of *Drosophila melanogaster* (Fig. 1a) have been maintained as large, isolated populations under distinct laboratory conditions^50,51^ for more than 1000 generations prior to the initiation of our experiment^50,52^ (Methods). As a result, these lineages exhibit substantial genomic divergence and partial, unidirectional, pre- and post-mating reproductive isolation. Most genetic variation is partitioned between the IV and LH_M_ lineages rather than within them (Fig. 1b) and absolute genomic divergence is contextually high (Fig. 1c). This level of divergence is comparable to what has been observed among geographically distant populations of *D. melanogaster*^53^, although it is lower than that typically found between recognized sister species within Drosophilidae^54–56^. Females from the IV population show higher mating rates with IV males than with LH_M_ males in no-choice assays (binomial GLM, χ^2^(1) = 12.49, *p* < 0.001; Fig. 1d) and mating occurs more rapidly with IV males than with LH_M_ males (Cox proportional hazards model, HR = 2.50, χ^2^(1) = 16.07, *p* < 0.001; Fig. 1e). IV females also produce roughly twice as many progeny when singly mated to IV males compared to LH_M_ males (LM, *F*_1,12_ = 9.803, *p* = 0.009; Fig. 1f). When IV females are sequentially mated to both types of males, IV males sire more than half of the offspring when they are first to mate, and nearly complete exclusion of LH_M_ paternity is observed when IV males are second to mate, consistent with conspecific sperm precedence^57^ (Fig. 1f).

**Figure 1.**
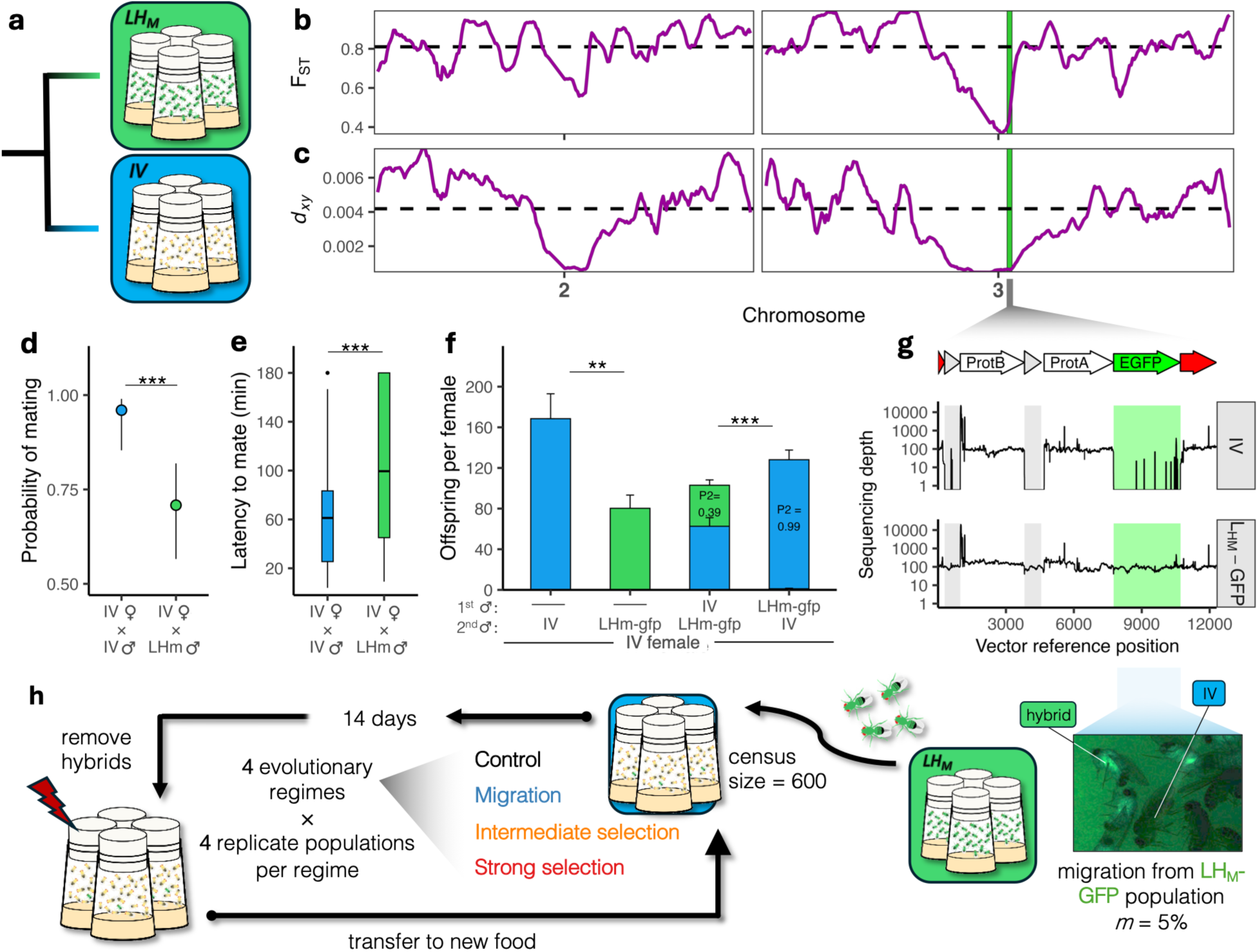
Genomic divergence and partial reproductive isolation between laboratory lineages enable an experimental test of reinforcement. (a) The LHM and IV laboratory lineages of *D. melanogaster* have been isolated from one another for minimally 45 years (>1000 generations)^50,52^. (b) Relative and (c) absolute genetic divergence of the lineages calculated in 2 Mb sliding windows with a 200 kb step size along the major *D. melanogaster* autosomes. Dashed lines denote autosomal mean. The vertical green line represents the location of the GFP insertion. (d) Probability of mating within a 3h period in a no-choice assay. (e) Latency to mating in the same assay, censoring pairs not mated within the 3h window. (f) Offspring production for singly- and doubly-mated IV females. Single bars (left) represent IV females singly-mated to each male type and stacked bars (right) represent IV females mated to both male types. (g) Whole-genome sequencing reads from IV and LHM-GFP mapped to the GFP insertion vector sequence. The shaded green region confirms the absence of the EGFP coding sequence in the IV genetic background. The presence of the insertion results in flies showing ubiquitous expression of EGFP. (h) Schematic of the experimental evolution design showing the four evolutionary regimes. ** *p* < .01 *** *p* < .001.

The presence of both pre- and post-copulatory reproductive isolating barriers between IV and LH_M_, together with substantial genomic divergence and standing genetic variation in IV (genome-wide mean *π* = 0.001; Extended Data Fig. 1), provides a uniquely powerful system to test whether reinforcing selection can strengthen reproductive isolation in the face of ongoing gene flow and to trace its genomic signature in real time. We imposed controlled secondary contact while manipulating the fitness consequences of hybridization. LH_M_ migrants (hereafter migrant males) were allowed to immigrate into replicate experimental populations founded from IV, replacing 5% of the resident adults each generation (Fig. 1h). Hybrids were identified via a dominant green fluorescent protein (GFP) insertion fixed in the LH_M_ population (Fig. 1h), allowing precise manipulation of postzygotic selection. We established two reinforcing selection regimes: an intermediate selection regime (I), in which 50% of hybrids were removed each generation, and a strong selection regime (S), in which 90% of hybrids were removed (Fig. 1h). In these regimes, the GFP^+^ allele acts as a classic postzygotic barrier locus imposing a fitness cost on females that produce hybrid offspring, thereby generating indirect selection favoring hybridization avoidance. To separate the effects of migration and selection, we also established a migration-only regime (M) which received migrants but experienced no experimentally imposed selection against hybrids, and an allopatric control regime (C), which experienced no migration and therefore no reinforcing selection. By holding the migration rate constant in treatments receiving migrants while manipulating the fitness cost of hybridization, the design allowed us to test how the strength of reinforcing selection influences the evolution of reproductive isolation in parallel replicated populations.

### The cost of hybridization governs the evolution of context-dependent reproductive isolation

Reproductive isolation can evolve during reinforcement through the strengthening of assortative mating or through postcopulatory mechanisms such as biased fertilization toward conspecific sperm. To capture both of these components, we measured total reproductive isolation in our populations after 114, 116, 119, and 130 generations of experimental evolution using an assay that integrates different stages of mating and fertilization and mirrors the social conditions under which flies evolved (Fig. 1h). Groups of virgin females from each evolved population were initially placed with evolved males from their own population, as occurs when a new generation of flies emerges in our evolutionary regimes, allowing initial within-population mating before the arrival of migrant males. These within-population groups were then transferred to new vials housing migrant males and allowed to interact for 3 days. All adults were then discarded and the paternity share of migrant males was recorded. Across all evolutionary regimes, hybridization rates were low (Fig. 2a), consistent with ancestral IV female avoidance of migrant males and the fact that most females likely had already mated to within-population males prior to migrant male arrival. Nevertheless, evolutionary regime had a strong effect on hybridization rate (evolutionary regime: χ^2^(3) = 12.80, *p* = 0.004). Compared to the control (C), hybridization was substantially reduced in the intermediate reinforcing selection (I) regime (OR_I-C_ = 0.438, *p* < 0.001) but not the strong reinforcing selection (S) regime (OR_S-C_ = 0.824, *p* = 0.403), or migration-only regime (OR_M-C_ = 1.309, *p* = 0.236) (Fig. 2a; per-generation: Extended Data Fig. 2).

**Figure 2.**
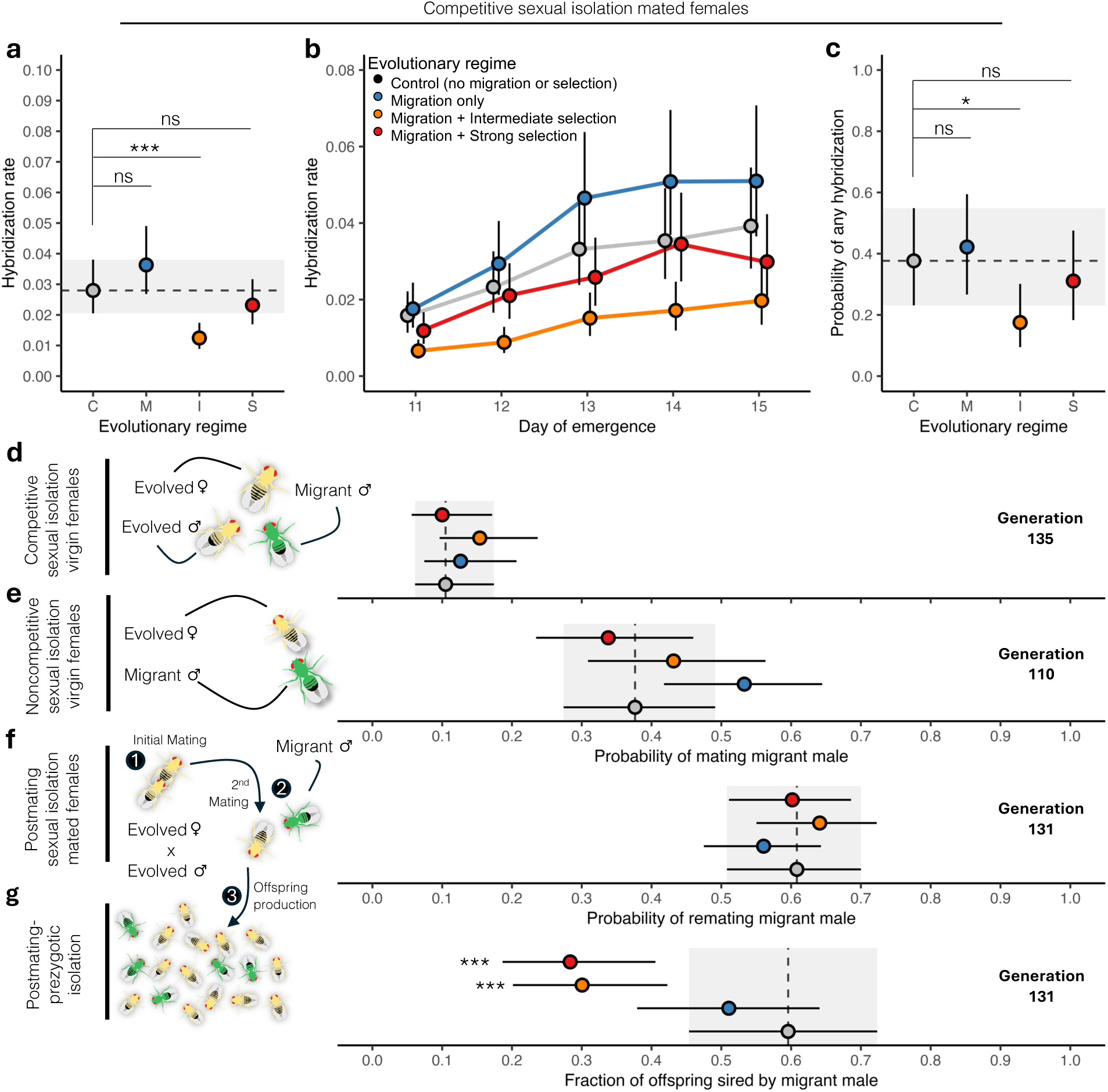
Reproductive isolation evolves after long-term experimental selection against hybridization. (a) Total reproductive isolation measured as the fraction of offspring sired by migrant males from groups of evolved females housed with evolved within-population males and migrant males, over five days of egg-laying. (b) Mean estimates of hybrid offspring production for each day of egg-laying. (c) The probability that a group produced any progeny fathered by a migrant male. a-c show the pooled analysis from measures obtained across 4 generations. (d) Competitive sexual isolation of evolved virgin females measured as the probability of mating a migrant male within 3h when simultaneously presented with an evolved within-population male. (e) Noncompetitive sexual isolation of evolved virgin females measured as the probability of mating a migrant male within 3h in a no-choice experiment. (f) Postmating sexual isolation of evolved mated females measured as the probability of remating a migrant male in a 24h period, 48h after mating to a within-population male. (g) Postmating prezygotic isolation of evolved females measured as the fraction of migrant male offspring sired by doubly-inseminated females when migrant male mates second. Points and lines represent evolutionary regime-wide marginal means ± 95% C.I. Asterisks indicate significant differences. * *p* < .05 ** *p* < .01 *** *p* < 0.001.

Offspring sired by migrant males tended to emerge later within the emergence window (Fig. 2b; day: χ^2^(4) = 533.154, *p* < 0.001), indicating that successful copulations by migrant males occurred after some delay. This pattern was consistent across regimes (evolutionary regime × day: χ^2^(12) = 14.021, *p* = 0.299). The probability that any migrant progeny emerged from a vial (i.e. that at least one female in the group mated with a migrant male) was lowest in the I regime and differed from the Control (OR_I-C_ = 0.438, *p* = 0.039), whereas the S and M regimes did not differ from Control (OR_S-C_ = 0.746, *p* = 0.875; OR_M-C_ = 1.208, *p* = 0.963), paralleling the reduced overall hybridization rate in I but not S or M (Fig. 2c). These results indicate that females from the I regime were less likely to mate with migrant males compared with females from the control, although lower fertilization success of migrant male sperm could also contribute. Regardless of the underlying mechanism, this result demonstrates that reinforcing selection can sharpen reproductive isolation under certain combinations of migration rates and selection against hybridization. The finding that reproductive isolation was strongest in populations experiencing intermediate levels of selection against hybridization highlights the dual role of migration during reinforcement: although gene flow erodes genetic differences between populations, maladaptive hybridization is necessary for selection to favor increased isolation^13,29,58,59^. Low migration rates limit the amount of maladaptive hybridization that can take place, reducing opportunities for reinforcement. Our results suggest that strong selection against hybridization may further limit these opportunities by reducing the recruitment of early-generation hybrids into the population and thus the potential for subsequent maladaptive backcrossing.

To determine which reproductive barriers responded to reinforcing selection, we partitioned total reproductive isolation into its premating and postmating components across the four evolutionary regimes over the course of 135 generations of experimental evolution. When virgin females were simultaneously presented with a within-population male and a migrant male, the within-population male won the overwhelming majority of contests, but this premating sexual isolation did not differ among evolutionary regimes at any generation measured (generation 135: Fig. 2d, earlier generations: Extended Data Fig. 3a). In no-choice assays, where virgin females encountered only a migrant male, females mated more often with migrant males but there was again no difference between regimes at any timepoint measured (generation 110: Fig. 2e, earlier generations: Extended Data Fig. 3b). This lack of divergence in virgin female mating behavior is perhaps unsurprising—in *D. melanogaster*, virgin females are highly receptive to mating, whereas mated females become less receptive due to physiological changes^60^, suggesting female choice may play a larger role during remating. Also, both female choice and male-male competition can occur through processes within the reproductive tract of multiply-mated females^61^. Nevertheless, premating sexual isolation is routinely found to have evolved between *Drosophila* species pairs^21,22,26^ and has been found in other experimental reinforcement work^36^. An alternative explanation for the lack of sexual isolation in virgin females is that it reflects the design of our evolutionary regimes, where females typically have already encountered within-population males before migrant males arrive, potentially limiting the scope for selection on premating preferences of virgin females. We therefore also examined postmating sexual isolation by testing the remating behavior of females previously mated to within-population males. After an initial mating, females were isolated for two days before being exposed to a migrant male for 24 hours. Although remating occurred readily within this interval (generation 131: Fig 2f, earlier generations: Extended Data Fig. 3c), no differences were observed between regimes. These results suggest that the reduced probability of any hybridization observed in the total reproductive isolation assay (Fig. 2c) depends on the simultaneous presence of both male types, consistent with an effect of the social environment on the evolution of reproductive isolation. In a higher density mating environment, females may encounter and evaluate many males in rapid succession, more closely matching the social conditions experienced during experimental evolution and more broadly in natural populations^62^. Finally, we measured postmating-prezygotic isolation by quantifying paternity of progeny from females first mated to within-population males and then to migrant males. Migrant males sired about half as many offspring when competing for I and S regime female reproduction as they did when competing for Control female reproduction (generation 131: OR_I-C_ = 0.292, *p =* 0.002; OR_S-C_ = 0.268, *p* = 0.001; Fig. 2g) and a similar number of offspring when competing for M regime female reproduction (generation 131: OR_M-C_ = 0.709, *p* = 0.389; Fig. 2g) (earlier generations: Extended Data Fig. 3d). This reduction in second-male success indicates that reproductive isolation strengthened in both I and S regimes through postcopulatory mechanisms, either via differences in copulation (e.g. duration or ejaculate transfer) or through processes within the female reproductive tract such as sperm competition or cryptic female choice.

The evolution of both postmating sexual isolation (Fig. 2c) and postmating-prezygotic isolation (Fig. 2g) in the I regime is consistent with patterns observed in natural *Drosophila* species pairs, in which premating and postmating prezygotic barriers evolve at comparable rates relative to postzygotic isolation^63^. However, we cannot determine the order in which these barriers arose in our experiment. The parallel responses of I and S in the postmating-prezygotic sexual isolation assay (Fig. 2g), compared with the response of I but not S in total reproductive isolation (Fig. 2a-c), suggest that the strength of reinforcing selection—and thus the cost of hybridization—differentially influences the evolution of specific isolating barriers. Because hybrids are recruited into I but rarely into S, females in I encounter more males carrying migrant ancestry earlier in the mating cycle, providing the opportunity for selection to favor mate choice barriers^64^. Females in S, by contrast, could initially mate with less risk of producing hybrid offspring. Once migrants arrive, however, biased fertilization toward sperm from within-population males would provide a large fitness benefit whenever remating does occur, even if such matings are infrequent^65,66^. Further work dissecting the mechanisms underlying these barriers may help test this interpretation. Intriguingly, we found no change in reproductive isolation of the M regime, relative to the Control, in any measures (Fig. 2) despite expectations that gene flow should erode reproductive barriers. This pattern raises the possibility that weak reinforcement also occurred in M, with baseline selection against hybridization offsetting the effects of gene flow.

### Selection against hybridization restricts introgression near the GFP barrier locus

To determine how reinforcing selection shaped genome-wide patterns of introgression, we sequenced pooled samples of female flies from the ancestral IV and LH_M_-GFP migrant populations prior to secondary contact and then from all evolved populations, including the migrant source population, after 100 generations of experimental evolution. Mapping of the GFP insertion revealed that the artificial postzygotic barrier locus resides at a TTAA landing site at position 4,474,594 on the right arm of chromosome 3 (Fig. 1g; Methods). Theory predicts that when much of the fitness cost associated with hybridization is concentrated at a single locus, selection should primarily restrict introgression at that locus and nearby linked genomic regions, while gene flow proceeds more freely elsewhere in the genome^32,67–69^. In hybrids that escape our experimentally imposed selection, independent assortment allows migrant alleles on the X and second chromosomes to be inherited without the barrier allele on chromosome 3. Crossing over in females more slowly uncouples migrant alleles on chromosome 3 from the barrier locus. Consequently, migrant alleles unlinked to the barrier locus are expected to introgress readily, whereas tightly linked alleles should remain associated with its deleterious effects and resist introgression.

To test these predictions, we estimated migrant ancestry (*α*; Methods) using 314,681 diagnostic sites fixed for alternative alleles between ancestral IV and LH_M_-GFP migrant populations (Fig. 3). Genome-wide, migrant ancestry differed among evolutionary regimes (LM, evolutionary regime, *F*_2,27_ = 21.36, *p* < 0.0001), with the highest levels in M (0.257 ± 0.011 SE), intermediate levels in I (0.210 ± 0.011 SE) and the lowest levels in S (0.153 ± 0.011 SE) (Extended Data Fig. 4). A principal component analysis (PCA) of allele frequencies at 743,566 SNPs showed a concordant pattern of regime-wise genomic separation from ancestral IV (Extended Data Fig. 5). Notably, the regime-level differences varied markedly among chromosomes (regime x chromosome interaction, *F*_4,27_ = 3.5, *p* = 0.02). Migrant ancestry was high and relatively similar among regimes on chromosome 2 (Fig. 3a). By contrast, chromosome 3, which harbors the GFP barrier locus, showed a stepwise increase in migrant ancestry: S populations had the lowest ancestry (0.0339 ± 0.02), I populations were intermediate (0.116 ± 0.02), and M populations had the highest ancestry (0.192 ± 0.02; Fig. 3b). The X chromosome showed a similar, though less consistent, regime-dependent pattern despite lacking the barrier locus (Fig. 3c). Overall, migrant ancestry was substantially lower on chromosome 3 and the X chromosome than on chromosome 2 (chromosome effect: LM, *F*_2,27_ = 188.7, *p* < 0.0001). Reduced X-linked introgression likely reflects the lower influx of migrant X chromosomes under male-only migration in our design; stronger selection on hemizygous X-linked variation may also contribute. These same patterns were apparent in chromosome-level PCAs (Extended Data Fig. 6).

**Figure 3.**
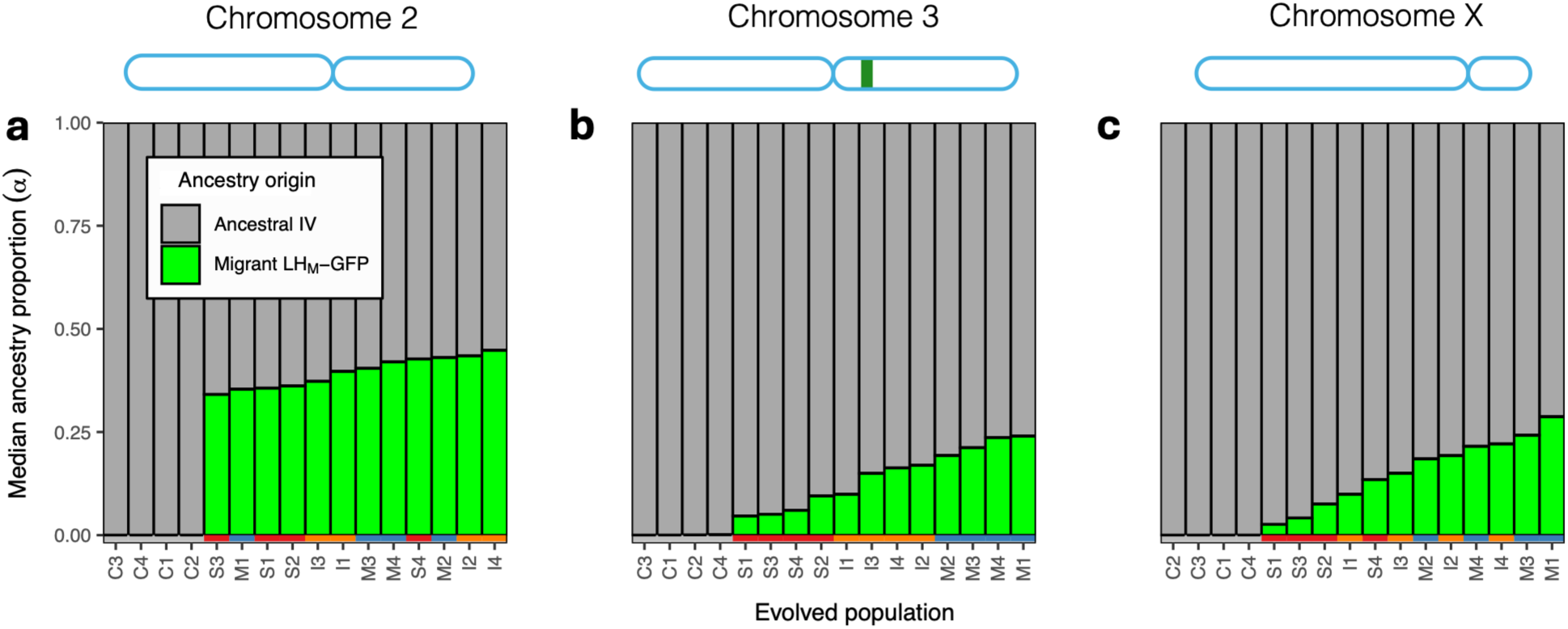
Chromosomal patterns of gene flow after 100 generations of continuous migration and selection against hybridization. (a-c) The contribution of ancestral population ancestry (*α*) to each of the 16 evolved populations per chromosome. (a) Chromosome 2, (b) Chromosome 3, with artificial barrier locus indicated in green, and (c) X Chromosome. Calculated from sites showing fixed differences between the ancestral IV and migrant populations.

We also estimated the per-generation effective migration rate^70^ (*m_e;_* Methods) in sliding windows across the genome, again using only sites fixed for different alleles in the ancestral IV and migrant LH_M_-GFP populations (including all sites did not change the overall pattern; Extended Data Fig. 7). As predicted for a localized barrier to gene flow, *m_e_* increased with distance from the barrier locus (Fig. 4a,b) and this increase was not explained by reduction in recombination rate near centromeres (Extended Data Fig. 8). The relationship between *m_e_* and distance from the barrier locus differed among regimes (distance × regime: *F*_2,6465_ = 168.21, *p* < 0.001), with M and I regimes showing an approximately fourfold steeper increase in *m_e_* with distance compared to the S regime (pairwise contrasts M-S and I-S *p* < 0.001; Fig. 4b). Compared to M, migrant ancestry *α* was reduced in I and S across a several megabase region flanking the barrier locus (Fig. 4c). Migrant ancestry *α* measured from 11 SNPs within 10kb of the barrier locus was reduced by 73.9% in I (t_9_ = 3.146, *p* = 0.028) and 65.2% in S (t_9_ = 2.776, *p* = 0.051) relative to M (Fig. 4d). All regimes, including M, showed significantly lower LH_M_-GFP introgression within this 10-kb region than the chromosome 3 average (*ᾱ*_chr3_ = 0.143; Fig. 4d), indicating that the GFP locus or closely-linked genetic variation carries a baseline fitness cost that is further elevated in I and S due to experimental selection against hybrids.

**Figure 4.**
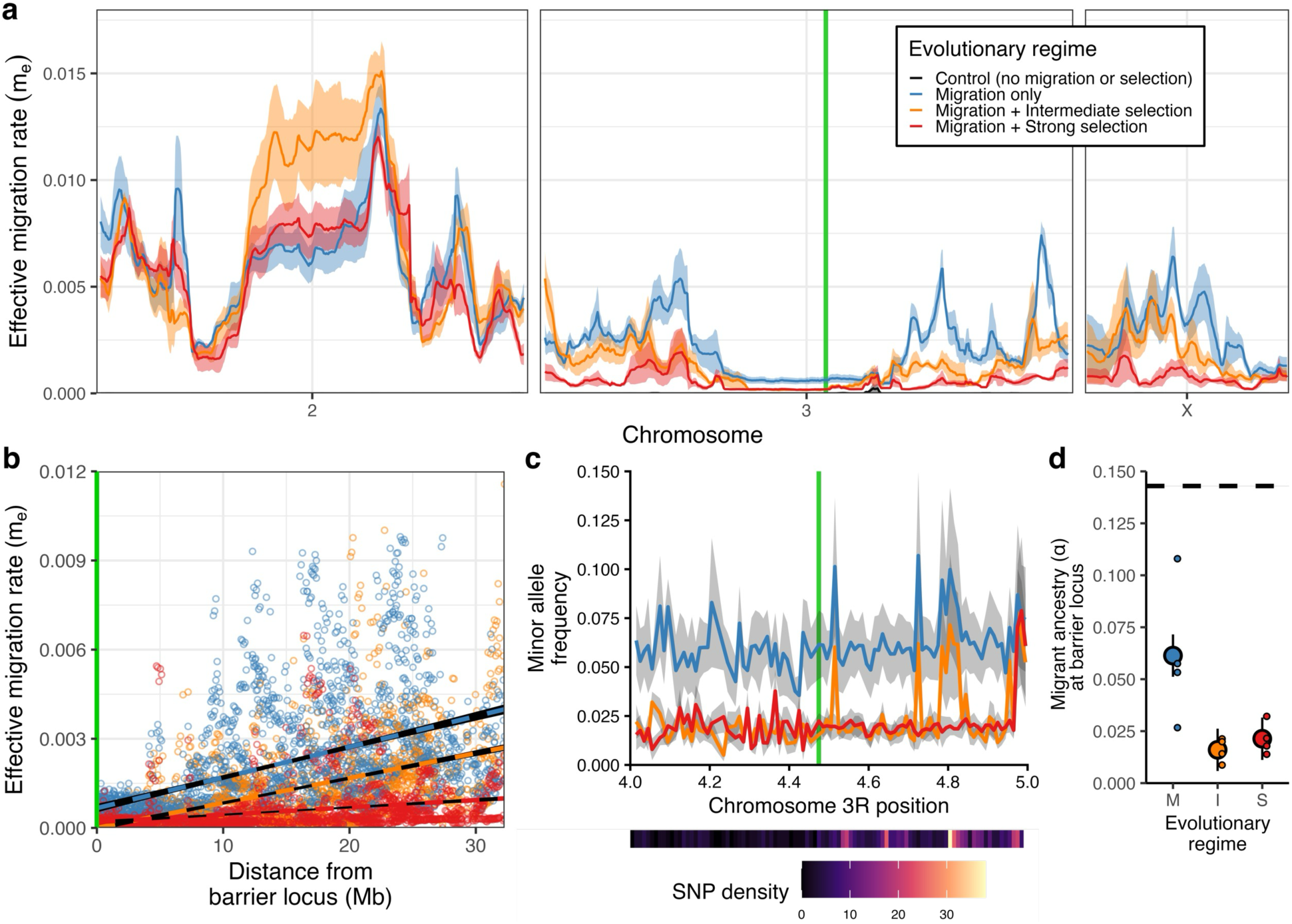
Selection against hybrids restricts introgression near the GFP postzygotic barrier locus. (a) Effective migration rate (*me*) calculated in overlapping 1Mb windows with a 100kb step size plotted across all major chromosomes. Lines show evolutionary regime-level means and shaded area shows ± 1 SE. (b) Effective migration rate estimated in 100kb non-overlapping windows plotted as distance from the barrier locus. (c) Percent migrant ancestry *α* calculated in 10kb windows plotted across a 1Mb region containing the GFP+ marker. (d) Percent migrant ancestry *α* averaged from 11 SNPs in closest proximity to the barrier locus. Large points represent evolutionary regime-level marginal means ± 1 SE and small points represent population-level means. Dotted black line represents chromosome 3 average in panels c and d.

Together, these patterns indicate that the genomic consequences of gene flow differ markedly among autosomes, reflecting variation in linkage to the barrier locus. The high degree of introgression and greater homogenization of chromosome 2 across regimes suggests that regions unlinked to the barrier locus experience relatively free gene flow. In contrast, the structured and regime-dependent pattern observed on chromosome 3 supports the prediction that selection against hybridization restricts gene flow primarily at linked genomic regions while permitting extensive introgression elsewhere in the genome.

### Signatures of adaptive evolution are concentrated near the GFP barrier locus

Reinforcing selection could strengthen reproductive isolation by favoring genetic variants that reduce the probability of hybridization (Fig. 5). We therefore performed a genome-wide scan designed to identify allele-frequency changes that were inconsistent with expectations under gene flow alone (e.g. Fig. 5a). We identified SNPs where the I or S regimes differed significantly from C and showed allele-frequency changes in the opposite direction of gene flow, a pattern consistent with divergent selection (Fig. 5b). We also tested for this pattern in M, where it was expected to be rare due to the absence of experimental selection against hybrids (Fig. 5c). Of the 158,575 sites fixed in the migrant population but with standing variation in the ancestral IV, a total of 3,348 sites showed a signature of divergent selection in I, S, or M, at an FDR of 5% (Methods). This signature occurred at only 107 SNPs in M (0.07% of all tested SNPs), compared with 1,648 SNPs in I (1.04%) and 2,448 in S (1.54%). After excluding SNPs with this signature in M, 3,253 candidates remained across I and S: 811 were exclusive to I, 1,625 were exclusive to S, and 817 (25.1% of candidate SNPs) showed parallel responses in both regimes (Extended Data Fig. 9). All three of these groupings were significantly enriched on chromosome 3 relative to the genomic background (Fisher’s exact tests, all FDR-adjusted *p* < 0.001; Supplementary Table 1). Among S-exclusive candidates, 62.2% resided on chromosome 3 (1,010/1,625; OR = 1.49), compared with 87.9% of I-exclusive candidates (713/811; OR = 6.61) (Fig. 5d). Shared I-S loci showed the strongest chromosomal bias, with an approximately 184-fold enrichment on chromosome 3 (813/817; 99.5% of all shared sites). A majority of the I- and S-exclusive SNPs on chromosome 3 that showed a signature of adaptive evolution were concentrated in correlated genomic clusters (Fig. 5e,f). By contrast, only 46 SNPs spanning three independent genomic peaks on chromosome 3 displayed evidence of divergent selection in M, with little overlap with I and S sites (Fig. 5g). Of the 817 shared candidate SNPs in I and S, 800 fell within a single 800-Kbp region on chromosome 3L (Fig. 5e-f, clusters S-8 & I-7; Supplementary Table 2), encompassing a total of 74 genes (40 protein-coding). Within this peak, SnpEff^71^ functional-effect annotation predicted 0 high-impact variants and 20 moderate-impact missense variants distributed across 11 unique genes. The top two genes (*CG10710*, *cmb*) harbored almost half (9) of the moderate-impact variants (Supplementary Table 3). Permutation-based gene ontology (GO) term enrichment of candidate genes (Methods) revealed no significant overrepresentation of biological processes defining divergent selection candidate gene lists, whether shared between contrasts or unique to contrasts (Supplementary Table 4). To summarize the magnitude of regime-level divergent change on chromosome 3 for each contrast, we calculated the mean allele-frequency shift from C, calculated from the SNP with the largest magnitude change in each cluster. M showed the weakest allele frequency shift from C (20.9%), followed by S (22.3%) and I (25.0%) (Extended Data Fig. 10a). These differences were driven primarily by the region harboring the shared cluster in I and S (Fig. 5e-f, purple points), which contained the most SNPs of all clusters and displayed the largest allele frequency change from C relative to I- or S-exclusive SNPs (29.3%, I-C; 34.3%, S-C; Extended Data Fig. 10b). Surprisingly, none of the 3,348 divergent-selection candidate loci identified in I or S differed from both C and M, which would have been expected if gene flow shifted all allele frequencies toward the migrant allele while selection simultaneously drove allele frequencies in I or S in the opposite direction. This result suggests some degree of selection against hybridization in the M selection regime and is consistent with the earlier observation of RI being maintained in the M populations.

**Figure 5.**
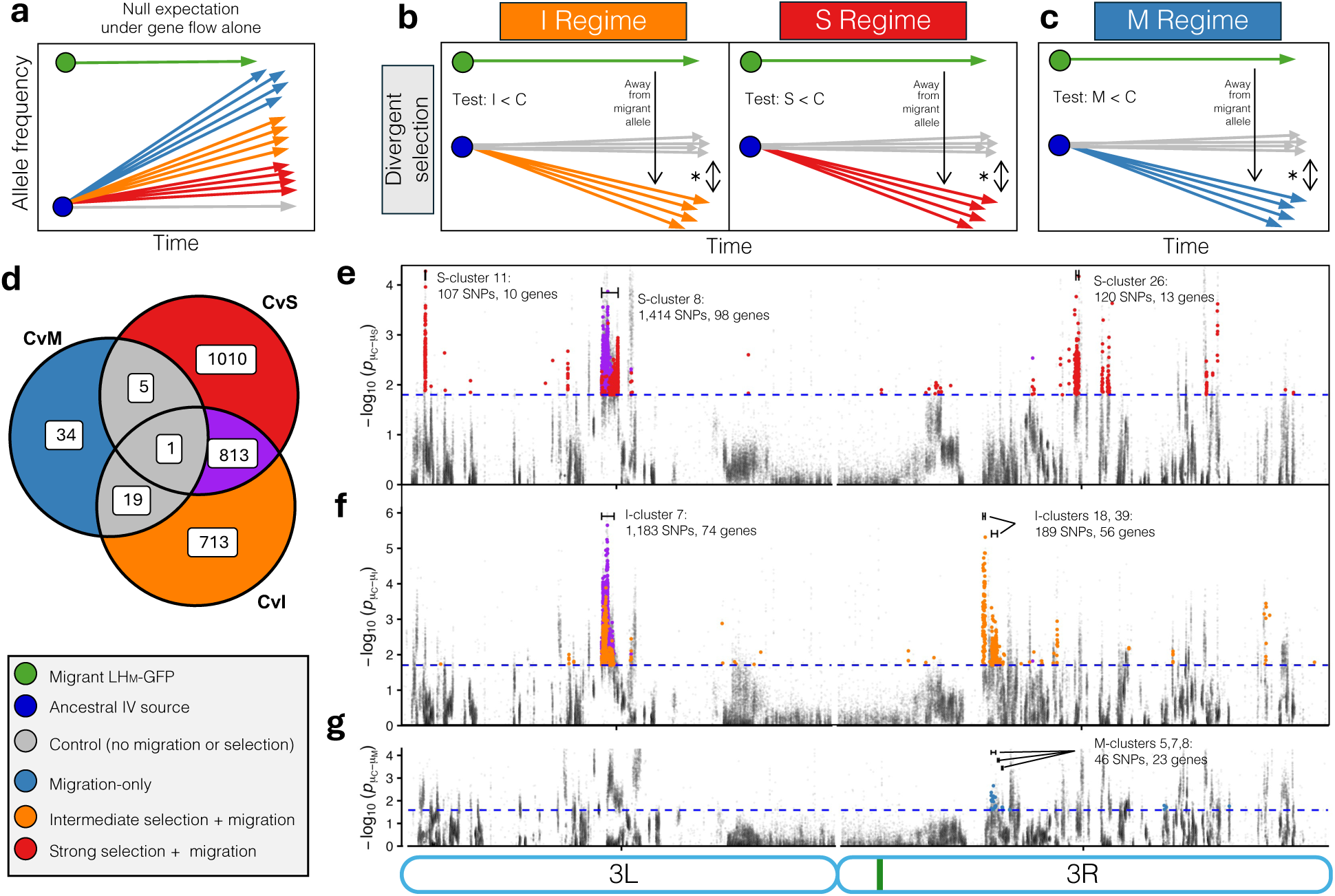
Patterns of adaptive allele frequency change enriched on the 3^rd^ chromosome. (a) Most of the genome should resemble a pattern driven primarily by gene flow. (b) SNPs with a signature of divergent selection were identified if the four replicate populations of the selection regimes (I or S) consistently differed from C in the predicted direction away from the migrant source population. (c) SNPs with a signature of divergent selection in the Migration-only (M) regime. (d) Venn diagram showing the overlap in chromosome 3 SNPs displaying a pattern of divergent selection of the M, I, and S contrasts. (e-g) Manhattan plots displaying the most significant divergent selection chromosome 3 SNPs in the (e) C-S, (f) C-I, and (g) C-M tests. Highly correlated SNPs were assigned to independently evolving clusters, with the three largest per contrast denoted by brackets. Dotted blue line denotes the FDR threshold of 0.05.

The strong concentration of divergent-selection candidates on chromosome 3 is notable given the location of the GFP barrier locus. Linkage to postzygotic barrier loci can facilitate both the establishment and the maintenance of variants contributing to reproductive isolation under ongoing gene flow^13,32,72–74^. These variants could influence female discrimination or, alternatively, affect male traits involved in mating and fertilization. For example, female discrimination against migrant-like phenotypes could favor further divergence of resident male traits. By limiting introgression of migrant alleles, selection at the GFP barrier locus would then facilitate this response to sexual selection in linked regions. Genomic signatures alone cannot distinguish changes in male traits from female responses, but the concentration of candidate loci within a few distinct genomic regions provides clear targets for future mapping of the causal loci underlying increased reproductive isolation.

## Conclusions

Together, our results provide a direct experimental demonstration that selection against hybridization can strengthen reproductive isolation between previously naïve populations despite ongoing gene flow. Evolved reproductive isolation was strongest under intermediate selection against hybrids. This pattern may reflect reduced hybrid recruitment under strong selection, limiting subsequent backcrossing and further opportunities for selection to favor avoidance of early-generation hybrids. The expression of premating reproductive isolation was also context dependent in our experiments, with reduced female remating with migrant males evident only when both resident and migrant males were present in the mating environment. Genomic divergence among regimes was concentrated near the barrier locus, while much of the remainder of the genome showed substantial introgression. Signatures consistent with adaptive evolution were enriched on chromosome 3, with many correlated candidate SNPs clustered in several distinct genomic peaks flanking the barrier locus. Collectively, these findings support a central prediction of reinforcement theory: selection against hybridization can increase reproductive isolation without requiring genome-wide barriers to gene flow, lending support to its long-proposed role in promoting speciation.

## Acknowledgments

We would like to thank Skylar McNulty for assistance with measuring ancestral population mating phenotypes and Teaghan Brosokas for help with DNA extractions and preparation of DNA pools for sequencing.

## Author contributions

Conceptualization: BH, GCM. Methodology: BH, GCM. Investigation: GCM, BH, SS, VB. Visualization: GCM. Funding acquisition: BH, GCM. Project administration: BH. Supervision: BH. Writing – original draft: GCM, BH. Writing – review & editing: GCM, BH.

## Data, code, and materials availability

Raw sequencing data will be available via the NCBI Sequence Read Archive under BioProject PRJNA1400512 upon publication. All phenotypic data, processed genomic data, and scripts will be deposited on Dryad at https://doi.org/10.5061/dryad.7d7wm388x upon publication.

## Materials and Methods

### Fly populations and baseline measures of reproductive isolation

The IV population of *Drosophila melanogaster* was originally founded in 1975 from ∼200 flies collected by P.T. Ives in Amherst, Massachusetts^50^. IV has since been maintained as a large, outbred population on a 14-day transfer schedule and shows high levels of genetic diversity alongside a clear signature of lab adaptation^50^. The LH_M_ population was founded in 1991 by Larry Harshman from ∼400 inseminated females collected in Modesto, California^51,52^ and is also outbred and maintained at a large census size, with non-overlapping generations. The IV and LH_M_ populations had therefore been isolated from one another for 1200+ generations at the start of this experiment. This isolation, combined with unique selective environments owing to differences in historical maintenance^50,51^, provided ample opportunity for some level of reproductive isolation to evolve. To enable identification of hybrids between IV and LH_M_ adults, we used a transgenic GFP-marked population that was derived from the LH_M_ population. This population, called LH_M_-GFP, was created in 2010 by backcrossing an LH_M_ mutant *pBac{Ubnls-EGFP, ProtB-EGFP}* into the ancestral LH_M_ genetic background and therefore involved some degree of bottleneck. The protB-GFP transgenic line is fixed for a dominant allele driving expression of GFP wherever ubiquitin is expressed (almost all tissue of the fly) while simultaneously driving expression of GFP to the head of the sperm^75^.

We measured baseline levels of assortative mating and sperm precedence between the IV and LH_M_-GFP populations with a no-choice experiment, presenting 3d old virgin IV females with 3d old virgin IV or LH_M_-GFP males. We recorded whether or not mating occurred within 3h and, when it did, the latency to mate. Immediately upon completion of mating, we removed the male and isolated the female for 48h. A subset of singly-mated females were then moved to a new vial while the remaining females were presented with a 7d old virgin male from the population not yet encountered and observed until mating. For matings that did not occur during the observation period, we allowed 24h of total interaction and successful second-male mating was inferred by the presence of either fluorescing or wild-type progeny. Immediately following mating, or after 24h with the male, the female was moved to a fresh food vial and allowed to oviposit for 4d before removal. Females that were not presented a second male were handled in the same way in order to obtain within-type progeny production measures from singly-mated females. The number and paternity of all offspring were scored 15d after females were moved to this final vial to measure within- and across-population progeny production and sperm precedence.

### Experimental evolution design

At the start of experimental evolution, populations were arbitrarily assigned to either a control regime (C), a migration-only regime (M), an intermediate selection + migration regime (I), or a strong selection + migration regime (S). Control populations were founded from 600 adult IV flies, while all other populations were founded from 570 adult IV flies and 30 migrants, as outlined below. This process resulted in 4 replicate populations of each regime (16 total populations: C1-C4, M1-M4, I1-I4, and S1-S4). Every 14 days, a set of four new bottles was established for each population, with 150 flies in each bottle, maintaining the census size at 600 individuals. In each C population, a sample of adults from the four bottles was used to establish four new bottles, each with 150 individuals. In M, I, and S populations, a sample of the flies occupying the four bottles was observed under fluorescent light and scored for the expression of green fluorescent protein. To mimic varying degrees of reinforcing selection, a fraction of the total number of GFP^+^ flies were removed each generation as prescribed by the selection regime: 0% in M, 50% in I, and 90% in S. Then, four new bottles were established, using 570 individuals from the sample and 30 new migrant males from the LH_M_-GFP source population, resulting in 150 individuals per bottle as in the C populations. In this way, the migration rate into M, I, and S populations was 5% and all populations in all evolutionary regimes had a census size of 600 individuals every generation.

### Measuring reproductive isolation in evolved populations

At various timepoints spanning 135 generations of experimental evolution we measured reproductive isolating barriers that occur before mating (premating barriers), and barriers that occur after mating but before fertilization (postmating-prezygotic isolation). Sexual isolation was separately measured in virgin and mated females, providing both premating and postmating sexual isolation measures. Both forms of sexual isolation were also measured in competitive and noncompetitive social settings. In addition, total reproductive isolation was measured with an assay designed to integrate both pre- and post-mating prezygotic barriers that was modeled after the environment in which the fly populations evolved. For all assays, flies were randomly subsampled from each population and reared in common gardens on identically prepared food vials at densities of 20 flies per vial. 10-12d after establishment of these vials, groups of 20 virgin males and females were collected under CO_2_ anesthesia and aged for 3-4d. Competitive sexual isolation in virgin females was measured at generation 22, 23, 44, 45, 63, 64, 86, and 135. Here, focal females were randomly placed in a mating vial alongside both a within-population male and a migrant male and observed until mating occurred. Each copula was observed under fluorescent light and the winning male was scored for fluorescence. Non-competitive sexual isolation in virgin females was measured at generations 45, 64, 90, 91, 97, and 110. For these assays, focal females were randomly placed in a vial with a migrant male and scored for whether mating occurred within 3h. Sexual isolation of mated females was measured at generations 64, 92, and 131. For these assays, a virgin evolved female was placed with a virgin within-population male, observed until copula formation, and then isolated immediately after completion of mating for 2d. The female was then moved to a fresh food vial with a 7d old virgin migrant male. The pair was given 24h of interaction before the migrant male was removed. 10d later, all progeny from the vial were scored for the presence or absence of fluorescence to infer whether the female had remated. For females who successfully remated the migrant male, we further quantified postmating-prezygotic isolation by recording the number of GFP^+^ and wild-type progeny that had emerged by 15 days after females first encountered the migrant male. GFP^+^ and wild-type individuals were counted at both day 12 and 15 for generation 131 and only at day 15 for generations 64 and 92. Finally, the total reproductive isolation assay was conducted at generations 113, 116, 119, and 130. Here, 5 evolved females were placed with 5 within-population males immediately upon collection as virgins. Three days later, males and females were moved to a fresh food vial alongside 5 3d-old virgin migrant males. These 15 individuals were allowed to interact for another 4 days before being discarded. The number of GFP^+^ and wild-type flies emerging from these vials were counted every morning for 5 days beginning 10 days after establishment of the second vial.

### Statistical analysis of reproductive isolation phenotypes

All statistical analyses were conducted with R v4.4.1^106^. To test for assortative mating in the ancestral IV and LH_M_ lineages, generalized linear models were fit with mating status (mated or not mated) as a binomial response variable and male as a fixed effect using the base R glm() function followed by a test for significance via a likelihood-ratio test (LRT). We tested for differences in mating latency using a Cox proportional hazards model^76^, treating latency to mate as the response and censoring pairs for which mating did not occur by 180 minutes. Male population was modeled as a fixed effect and significance was assessed with a LRT implemented in the Survival package v3.3.3^77^. For reproductive output of singly mated females, we fit a linear model with total number of offspring as the response variable and male population as a fixed effect with the effect of first-male type evaluated using an F-test. Paternity share of doubly mated females was modeled using a generalized linear model with a logit-link function for the response variable of the proportion of IV or LH_M_ offspring with first male-type to mate as a fixed effect. The effect of mating order on paternity share was tested using a LRT.

For all assays involving evolved populations, we tested for a block effect between consecutive generations of the same measure, collapsing the data into a single generation if there was no effect. To assess the probability of mating or remating in the evolved populations in our competitive and non-competitive sexual isolation assays we used the mixed() function implemented in the afex package v1.4.1^78^, a wrapper for the lme4 package v1.1.37^79^. We fit a binomial generalized linear mixed-effects model (GLMM) of mating status (mated or remated coded as 1) as a function of evolutionary regime, generation, and their interaction, with replicate population as a random effect. For assessing the proportion of migrant male progeny in the postmating-prezygotic assay we again used the mixed() function to fit a binomial GLMM but here using a logit-link function with the number of migrant or within-population male offspring as the response variable and fixed effects of evolutionary regime, generation, and their interaction, with replicate population as a random effect. In the total reproductive isolation assay we used the same model to estimate the overall effect across all days and fit an additional model including a fixed effect of day and its interaction with evolutionary regime and a random effect of female. For all models, fixed-effect significance was evaluated using a nested model Type-III LRT. Evolutionary regime-level marginal means and pairwise mean differences were obtained with the emmeans package v1.11.2^80^, reported on the response scale with back-transformed (logit) 95% CIs unless otherwise noted.

### Pooled-sequencing bioinformatics pipeline

DNA was isolated from the ancestral IV stock used to establish the experimental populations, the LH_M_-GFP stock used to establish the migrant source stock, the migrant source stock at generation 100, and from each evolved population after 100 generations of experimental evolution. For each population, ten groups of 30 females were obtained and randomly divided into two replicate samples, each totalling 150 females. Flies were homogenized using the BeadBug3 tissue homogenizer with 70mm zirconium beads (Benchmark Scientific). DNA was extracted using the DNeasy Blood and Tissue kit (Qiagen) following manufacturer protocols. DNA purity was assessed using a NanoDrop Lite spectrophotometer (ThermoFisher Scientific) and concentrations were quantified using a Qubit 4 fluorometer with broad range dsDNA working solution (Invitrogen). Two equimolar DNA pools were constructed from 150 female flies each, resulting in 38 total pools. Paired-end 150 bp Illumina libraries were prepared for each pool and sequenced on an Illumina NovaSeq X instrument at the Duke University Sequencing and Genomic Technologies (SGT) resource. Raw sequencing data are available via the NCBI Sequence Read Archive under BioProject PRJNA1400512 and summarized in Supplementary Table 5. The two pools per population together represent 600 haploid chromosomes sequenced at a combined target depth of ∼120x, exceeding recommended thresholds for accurate allele-frequency estimation in Pool-seq data^81^.

Raw sequencing reads were quality trimmed with fastp v0.23.4^82^ and quality was assessed before and after trimming using FastQC v0.11.5^83^ and MultiQC v0.1.14^84^. Reads were then mapped to the *Drosophila melanogaster* reference assembly v109 (retrieved Oct. 2024 from Ensembl - NCBI Accession #GCA_000001215.4) using BWA-MEM v3.16^85,86^. Highest quality mate pairs were tagged with Samtools fixmate and duplicate reads were marked with Samtools markdup implemented in SAMtools v1.19.2^87^. Overlapping regions of paired-end reads were clipped using bam clipOverlap from bamUtil v1.0.15^88^. Reads flanking indels were realigned using GATK v3.8.0^89^. Variant calling was performed using bcftools mpileup followed by bcftools call to call variant positions. Reads with mapping quality <40 or base quality <30 were excluded during pileup generation, following recommended Pool-seq practices^90^. The resulting Variant Call Format (VCF) file was further subset using bcftools filter to retain only biallelic single nucleotide polymorphisms (SNPs) and to remove sites with an overall quality score (QUAL) of less than 20. Chromosome position, reference and alternative alleles and corresponding per-sample allele counts were extracted from the FORMAT/AD field using bcftools query and reshaped into a site-by-sample matrix using a custom python script.

A custom R script was used to polarize alleles, further filter alleles, and calculate allele frequencies. First, sites mapping to the Y chromosome were removed because all pools consisted exclusively of females. To reduce the influence of mis-mapped reads and highly repetitive sequences, sites with excessively high read depth defined as exceeding the 99th percentile of the depth distribution were excluded. Excessive depth thresholds were calculated separately for the core nuclear genome (chromosomes 2 and 3), the 4th “dot” chromosome, and the X chromosome following recommendations of Burny et al. (2020)^91^. Sites mapping to the mitochondrial genome and rDNA were excluded due to their abnormal coverage distributions. To ensure reliable baseline allele-frequency estimates for downstream analyses, sites were further filtered based on minimum read depth. Sites were retained only if all ancestral pools (IV and migrant LH_M_-GFP) and evolved populations had a minimum depth of 20 reads at that position for each sequencing replicate. Sites were also required to have a minimum allele count of 4 reads for each allele across all ancestral pools combined. This criterion ensured that sites were supported by sufficient ancestral variation while avoiding conditioning site inclusion based on allele frequencies observed in evolved populations, particularly sites that may have moved to fixation during experimental evolution and would therefore be monomorphic in the evolved populations. Finally, sites showing inconsistent allele frequency estimates among sequencing replicates, defined as >0.25 standard deviation from the mean estimate of the two replicates, were removed. Counts were then summed across both 150 individual sequencing replicates for each sample resulting in 19 population pools which were used for all downstream analyses. The final SNP dataset contained 743,070 SNPs with a mean per-sample sequencing depth of 129.58x (median 130.21x) that was fairly consistent across population pools (range: 114.81x-151.27x; Supplementary Table 6).

### Mapping the EGFP locus in the migrant LH_M_-GFP population

The EGFP marker fixed in the LH_M_-GFP population is the product of transposon-mediated transgenesis resulting in a random genomic landing site. To identify the location of the ubiquitous EGFP insertion (the experimentally-imposed postzygotic barrier locus) in the LH_M_-GFP genetic background, we first screened raw FastQ files of the ancestral LH_M_-GFP populations for reads containing EGFP-specific sequences looking for exact matches via grep, yielding a starting search point. We then performed iterative read-walking in both directions and periodically queried the growing contig by BLASTn^92^ to identify transitions from transgene sequence to *Drosophila melanogaster* genomic DNA, using FlyBase JBrowse (release FB2025_25, genome r6.66^93^) to confirm the identity of protein-coding regions and transitions to genomic DNA. Identification of the correct target site was further validated by the presence of TTAA tetranucleotide motifs flanking the insertion, which is the preferred landing site for the *PiggyBac* transposon^94^. This process ultimately assembled a ∼12 kb contiguous sequence spanning vector sequences, the *protA* and *protB* coding sequences, the *EGFP* coding sequence, and flanking genomic DNA on both sides. The contig’s composition was validated by BLASTn and FlyBase JBrowse (vector backbone, *EGFP* and *protA* and *protB*) and its genomic coordinates were assigned by best hits to the *D. melanogaster* reference. Finally, to verify the insertion, we aligned LH_M_-GFP and IV reads to this contig with BWA using default parameters^85^, inspected coverage and looked for split-read/discordant-pairs. Break-point support was evident due to zero coverage at the EGFP sequence in IV and high coverage in LH_M_-GFP, confirming presence of the insertion only in the LH_M_ background as well as the location of the insertion in the genome.

### Genetic diversity and genetic differentiation of ancestral populations

Within-population nucleotide diversity was quantified in the ancestral IV and LH_M_-GFP migrant populations as *π* (also known as Tajima’s *π*), a generalizable metric defined as the probability that two randomly chosen sequences differ at a randomly chosen locus. We applied the pool-seq–corrected *π* estimator *θ_π_* as implemented in grenedalf v0.6.2^95^, to calculate nucleotide diversity averaged over genomic windows including SNPs and invariant sites by inputting an all-sites table and specifying --averaging-policy valid-loci. This all-sites table included alleles summed across sequencing replicates including invariant sites and SNPs passing all filters outlined previously. Invariant sites with a total depth of less than 20 were set to missing. We further used this all-sites table to characterize the distribution of genetic variation between our ancestral populations. The fixation index F_ST_ namely F_ST_ *^Hudson^* ^96,97^ was applied using the pool-seq corrected estimators as implemented in grenedalf v0.6.2^95^. Specifically, F_ST_*^Hudson^* measures the reduction in expected heterozygosity *H_e_* due to population structure by comparing within-population sequence diversity *π*_within_ to between-population divergence *π*_between_ (aka *d_xy_*).

### Population genomics of evolved populations

#### Principal components analysis

To visualize genome-wide and chromosome-specific patterns of allele-frequency covariance among ancestral IV, migrant, and evolved populations, we performed principal component analysis (PCA) using the randomAllelePCA() function implemented in the R package poolfstat v3.0.0^98^. For each SNP and population, this approach randomly samples a single read, generating a population-by-SNP matrix for single value decomposition. By assigning one randomly sampled read to each population at each SNP, this method accounts for unequal sequencing coverage among pool-seq samples.

#### Per-generation effective migration rate

We estimated per-generation effective migration rate, here termed *m_e_*, using the estimator from Wang & Whitlock (2003, eq. 8)^70^. We removed windows where any population showed an *m_e_* estimate of 1. This can occur if the allele frequency of the recipient population moved to match the allele frequency of the migrant population in a single-generation-like pattern, usually from windows harboring a limited number of SNPs. Genomic windows where any of the control populations displayed *m_e_* values greater than 0.001 migrants per generation were excluded to reduce background noise that cannot be explained by gene flow.

#### Hybrid index (*H*) & Ancestry index (*α*)

To quantify patterns of introgression, we calculated the hybrid index, *H*, for each locus *i*, defined as the relative shift in allele frequency between the ancestral island population (generation 0) *q*_0_ and the evolved island population at timepoint *t*, *q_t_*, scaled by the initial allele frequency difference between the ancestral island population and the ancestral migrant population *q_m_* following eq. 8.16 (Hedrick, 1983^99^):

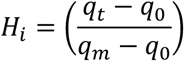

Sites where *q*_0_ = *q_m_* were filtered out as these are not informative for introgression. When we computed *H* from only fixed sites, we refer to *H* as the ancestry index *α*.

To test whether the degree of admixture was affected by evolutionary regime or chromosome, we fit a linear model using ordinary least squares using the lm() function in base R^106^ with median *α* values as the response variable for each level of population and chromosome as predictor variables. Estimated marginal means and pairwise contrasts were obtained from the fitted model using the emmeans package v1.11.2^80^. Marginal means were computed for the main effects of treatment and chromosome, as well as their interaction.

### Genome scan for adaptive evolution

We identified a signature of ‘divergent selection’ at sites for which either selection regime (I or S) or the migration-only regime (M) had shifted away from the migrant population relative to the control regime (C). We fit separate models contrasting C with I, C with S, and C with M. For each contrast, we fit a per-locus binomial generalized linear mixed model using allele counts as the response variable, evolutionary regime as a fixed effect, replicate population as a random effect, and sequencing pool nested within replicate population as an additional random effect. Significant loci were classified as showing divergent selection when the population receiving migration had a lower polarized allele frequency than C, indicating a shift away from the migrant-associated allele. For both models we excluded sites polymorphic in the migrant LH_M_-GFP population and sites where the initial allele frequency of IV had changed by more than 10% by generation 100 in C (i.e. drift- or error-prone sites). We assessed the significance of each regime effect with an LRT integrated into mixed(). False discovery rate (FDR) was calculated from the raw p-value distributions from the models using the Storey-Tibshirani method^100^, implemented in the qvalue R package^101^, and sites with q < 0.05 were retained as candidates.

### Defining correlated haplotype blocks

To avoid treating tightly linked candidate SNPs as independent observations^102^ and to summarize the genomic distribution of divergent evolutionary responses, we defined correlated haplotype blocks by adapting the residual-correction approach of Wyer et al. (2023)^103^ and Kawecki et al. (2021)^104^. For each contrast (CvI, CvS, and CvM), we obtained SNPs classified as candidates for divergent selection. At each SNP, we calculated allele-frequency residuals for each replicate population by subtracting the mean focal-allele frequency of its evolutionary regime from the observed focal-allele frequency in the population. We quantified the similarity of evolutionary responses between every pair of candidate SNPs as Pearson correlation coefficients (*r_w_*) between their residual allele-frequency vectors across populations. High positive correlation coefficients indicate that two SNPs deviate from their respective treatment in similar ways across replicate populations, consistent with hitchhiking via genetic linkage. We defined independent SNP clusters as correlated sets of candidate SNPs on the same chromosome that were separated by no more than 100 Kbp and had an *r_w_ >* 0.70. Specifically, we assigned SNPs to the same correlated haplotype block when they were linked directly, or indirectly through intervening SNPs, at the distance and correlation thresholds. We annotated candidate SNPs within each haplotype block using SnpEff^71^ and summarized the number of candidate SNPs and unique genes represented in each cluster. Functional effects were classified according to SnpEff impact categories based on how likely they are to impact phenotype (high, moderate, low, and modifier).

### Permutation-based gene ontology enrichment tests

To test whether particular gene ontology (GO) terms were overrepresented among genes containing candidate SNPs for each contrast (CvI, CvS, or CvM) while controlling for unequal SNP density among genes and non-independence among SNPs within genes, we conducted a permutation-based enrichment analysis mirroring the framework implemented in Gowinda^105^. All tested SNPs were mapped to annotated genes to define the respective genomic background for each category. Candidate SNPs were similarly mapped to genes, and multiple candidate SNPs within the same gene were collapsed such that each gene contributed to only a single observation. For each GO biological process (BP) term, we counted the number of candidate genes assigned to that term. To generate a null distribution, SNPs were sampled without replacement from the genomic background and mapped to genes until the number of unique sampled genes equaled the observed number of candidate genes. This procedure preserves differences among genes in their probability of being sampled due to variation in SNP density while treating all SNPs within a gene as a single linked unit. For each permutation, the number of sampled genes assigned to each GO term was recorded. Empirical p-values were calculated as the proportion of 10,000 permutations in which the number of randomly sampled genes annotated with a given GO term was at least as large as the number of candidate genes annotated with that term. Multiple testing was controlled using the q-value method, and GO terms were retained as candidates for enrichment at an FDR threshold of 0.05.

## Extended Data

**Extended Data Fig. 1:**
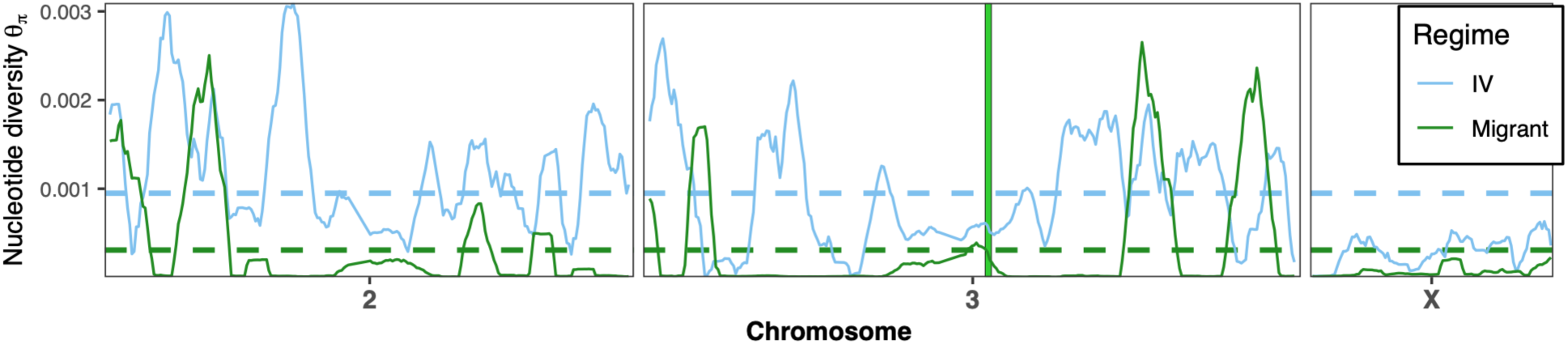
Genome-wide nucleotide diversity of ancestral IV and LH_M_-GFP. Nucleotide diversity of ancestral populations (IV: blue, LH_M_-GFP: green) computed across the major autosomes and sex chromosome in overlapping 2Mbp windows, with 200Kbp strides. Dotted lines represent genome-wide mean. The green vertical line designates the location of the GFP barrier locus.

**Extended Data Figure 2:**
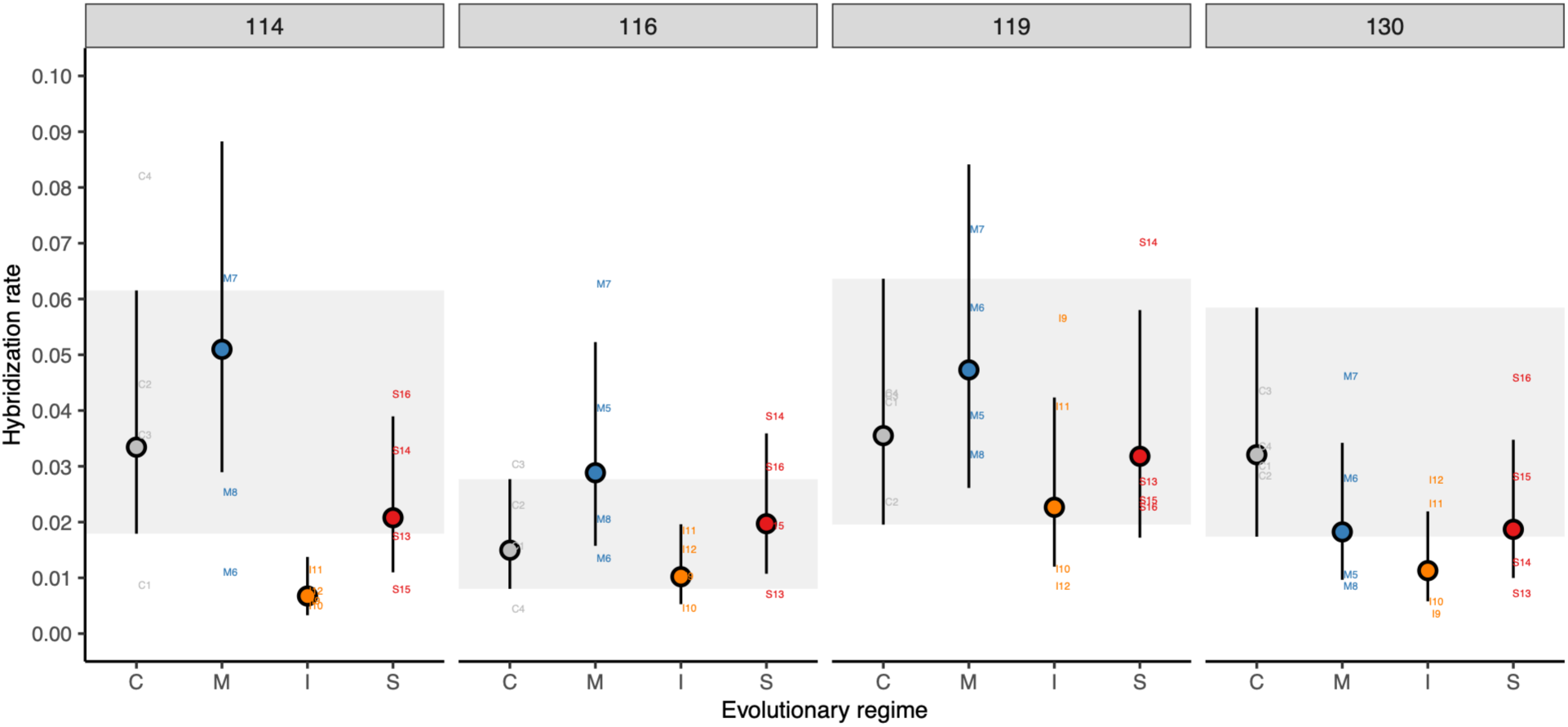
Hybridization rate in assays of total reproductive isolation across four generations. Facet labels denote generation measured. Points represent evolutionary-regime marginal means and text labels indicate replicate population-level averages.

**Extended Data Figure 3:**
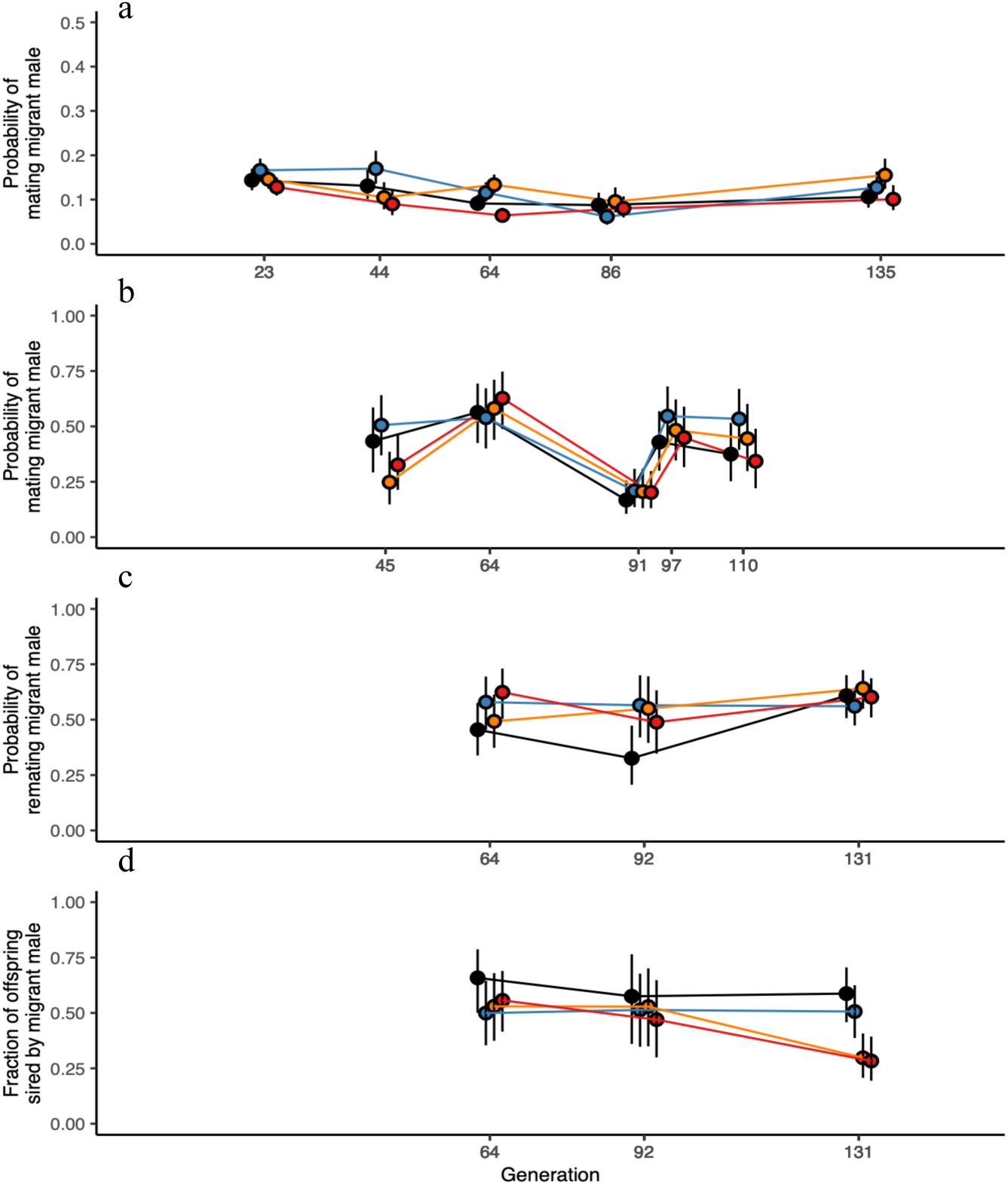
Measures of prezygotic reproductive isolating mechanisms over the course of 135 generations of experimental selection against hybridization. (a) Premating sexual isolation defined as the probability of a virgin evolved female mating a migrant male (b) Competitive remating sexual isolation defined as the probability of a virgin evolved female mating a migrant male when also presented with an evolved male. (c) Sexual isolation defined as the probability of an evolved female remating a migrant male after already having mated a within-population male. (d) Postmating-prezygotic isolation defined as the proportion of migrant offspring produced by an evolved female after having mated both male types. Colors correspond to evolutionary regimes: Control regime (black), migration regime (blue), intermediate selection regime (orange), strong selection regime (red).

**Extended Data Figure 4:**
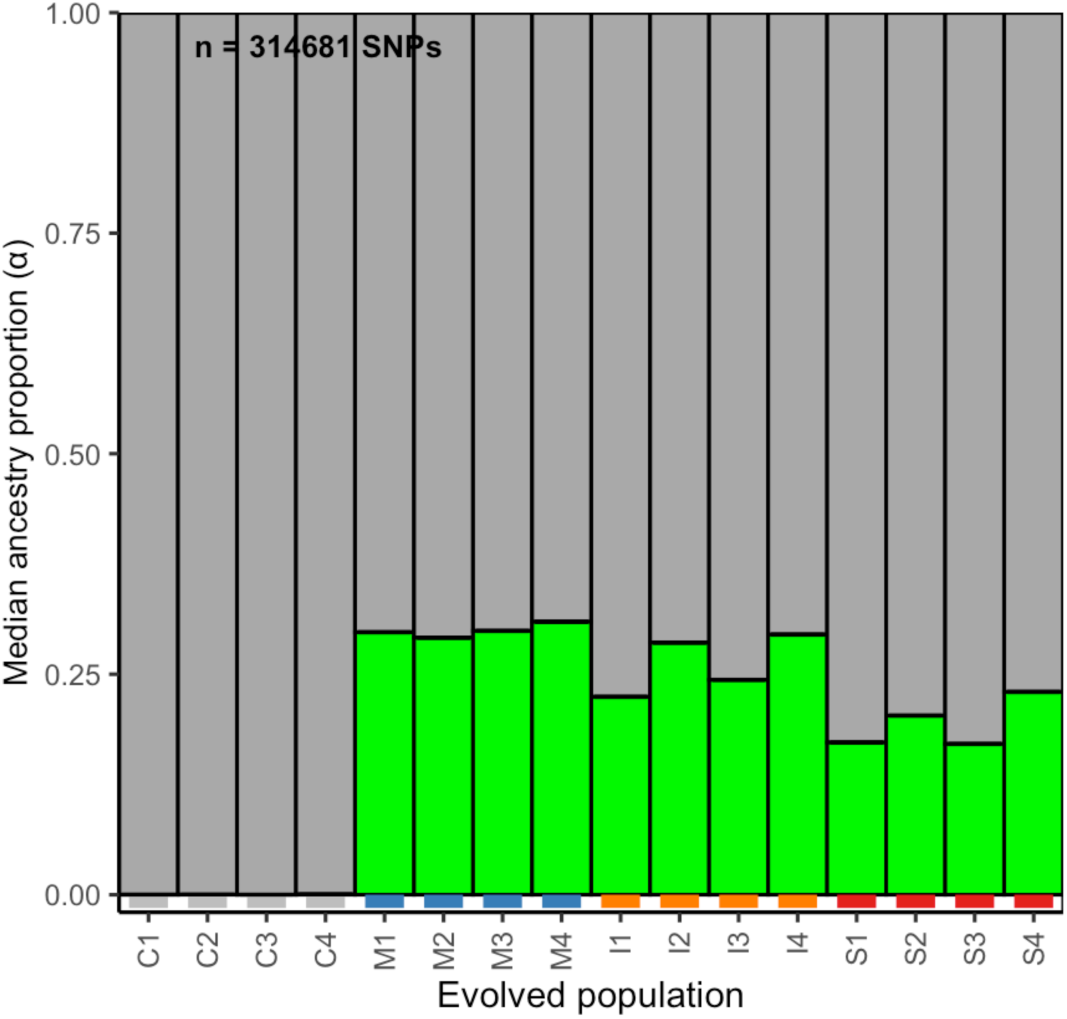
Median proportion of migrant ancestry per replicate population by generation 100. Averaged across 314,681 sites showing fixed differences between the ancestral IV and LHm-GFP spanning the entire *D. melanogaster* genome. Populations from the control regime (grey), migration regime (blue), intermediate selection regime (orange) and strong selection regime (red) are indicated

**Extended Data Figure 5:**
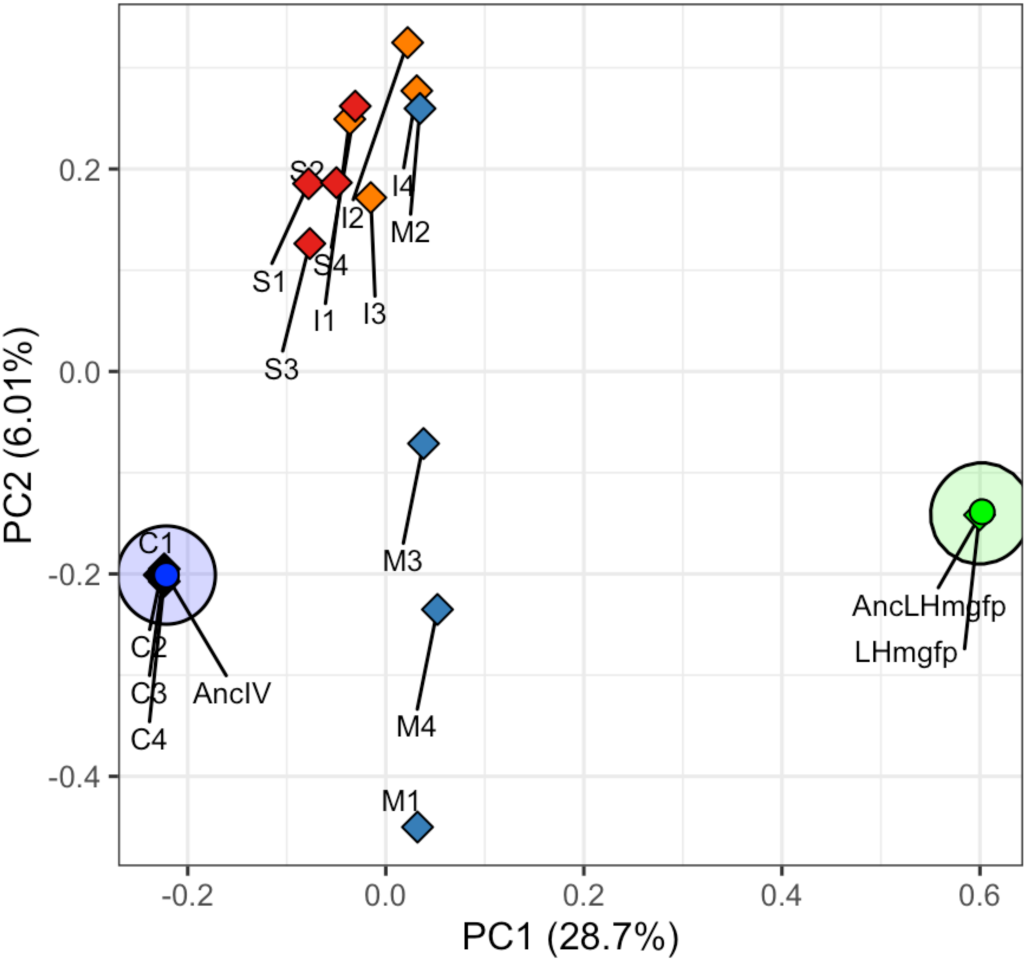
Genome-wide PCA of all 743,566 SNPs. Populations from the control regime (black), migration regime (blue), intermediate selection regime (orange) and strong selection regime (red) are indicated. The ancestral IV population is dark blue, while the ancestral and evolved LHM-GFP migrant populations are in green.

**Extended Data Figure 6.**
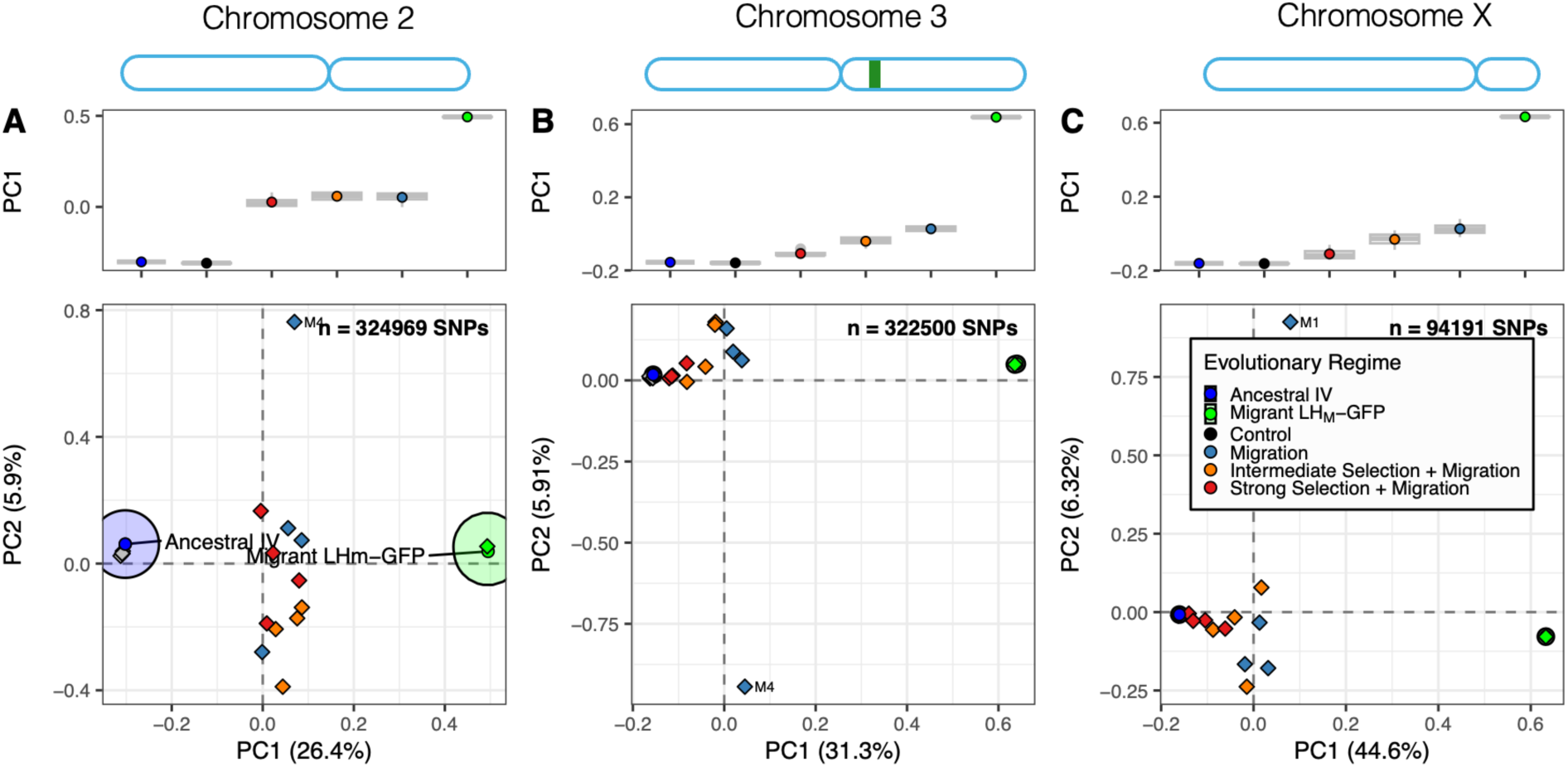
Principal components analyses of all SNPs from each major chromosome. (a) Chromosome 2, (b) Chromosome 3, (c) X chromosome. Each point represents a population. Populations from the control regime (black), migration regime (blue), intermediate selection regime (orange) and strong selection regime (red) are indicated.

**Extended Data Figure. S7.**
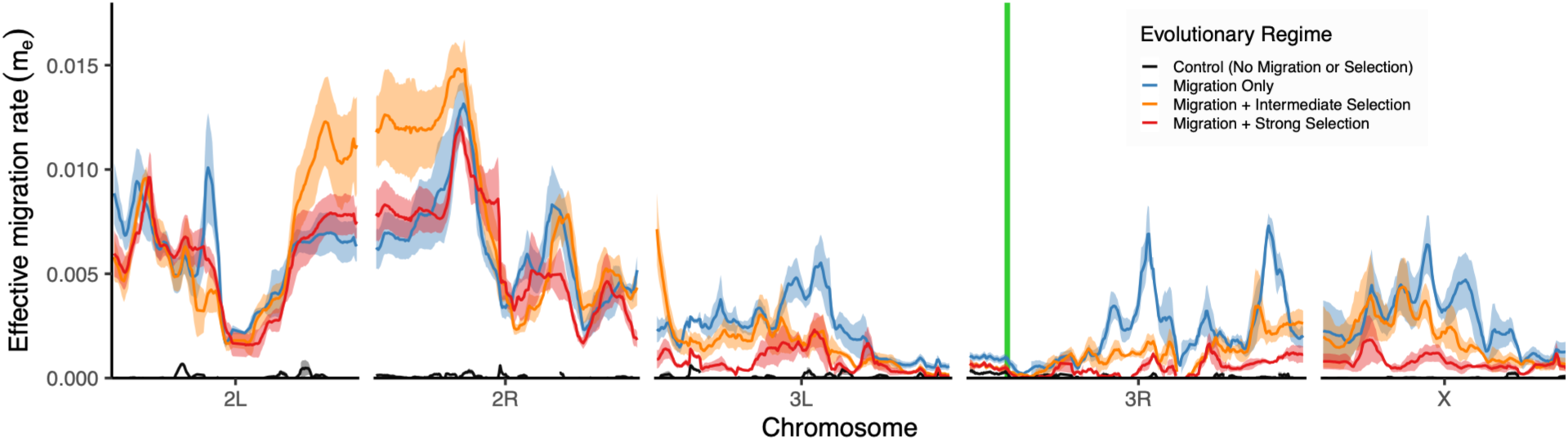
Effective migration rate in sliding windows including polymorphic and fixed SNPs. Populations from the control regime (black), migration regime (blue), intermediate selection regime (orange) and strong selection regime (red) are indicated. The location of the GFP barrier locus is drawn as a green vertical line. Windows containing a locus where any population showed an allele frequency increase greater than both the ancestral IV and ancestral LHM-GFP were filtered out as these result in a division by zero error in the *me* estimator and cannot be explained by gene flow.

**Extended Data Figure 8:**
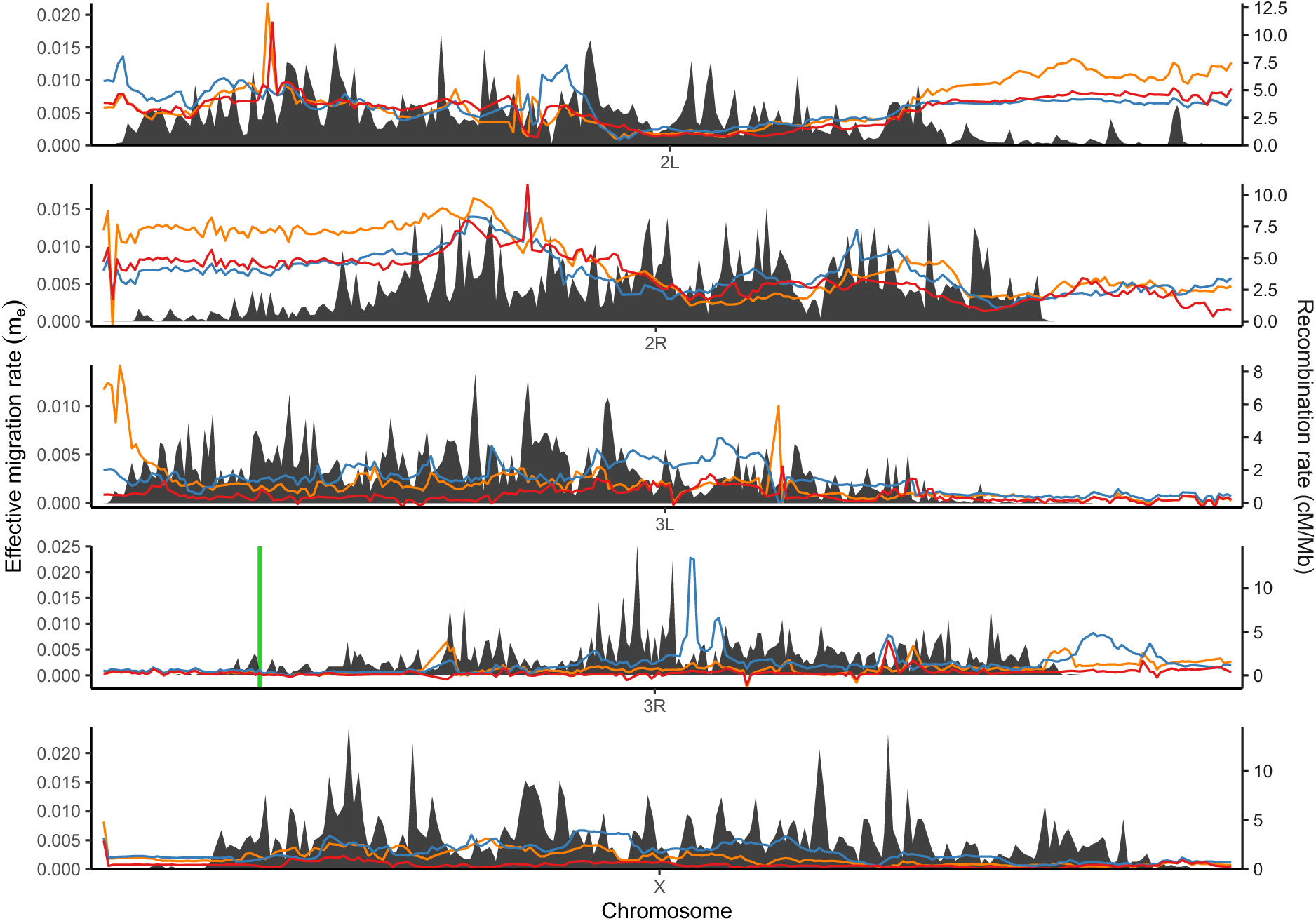
Effective migration rate plotted alongside recombination rate (crossover rate from ^1^) including only fixed sites. Populations from the control regime (black), migration regime (blue), intermediate selection regime (orange) and strong selection regime (red) are indicated as colored lines. The shaded region indicates the recombination rate and the green vertical line denotes the location of the GFP barrier locus.

**Extended Data Figure 9:**
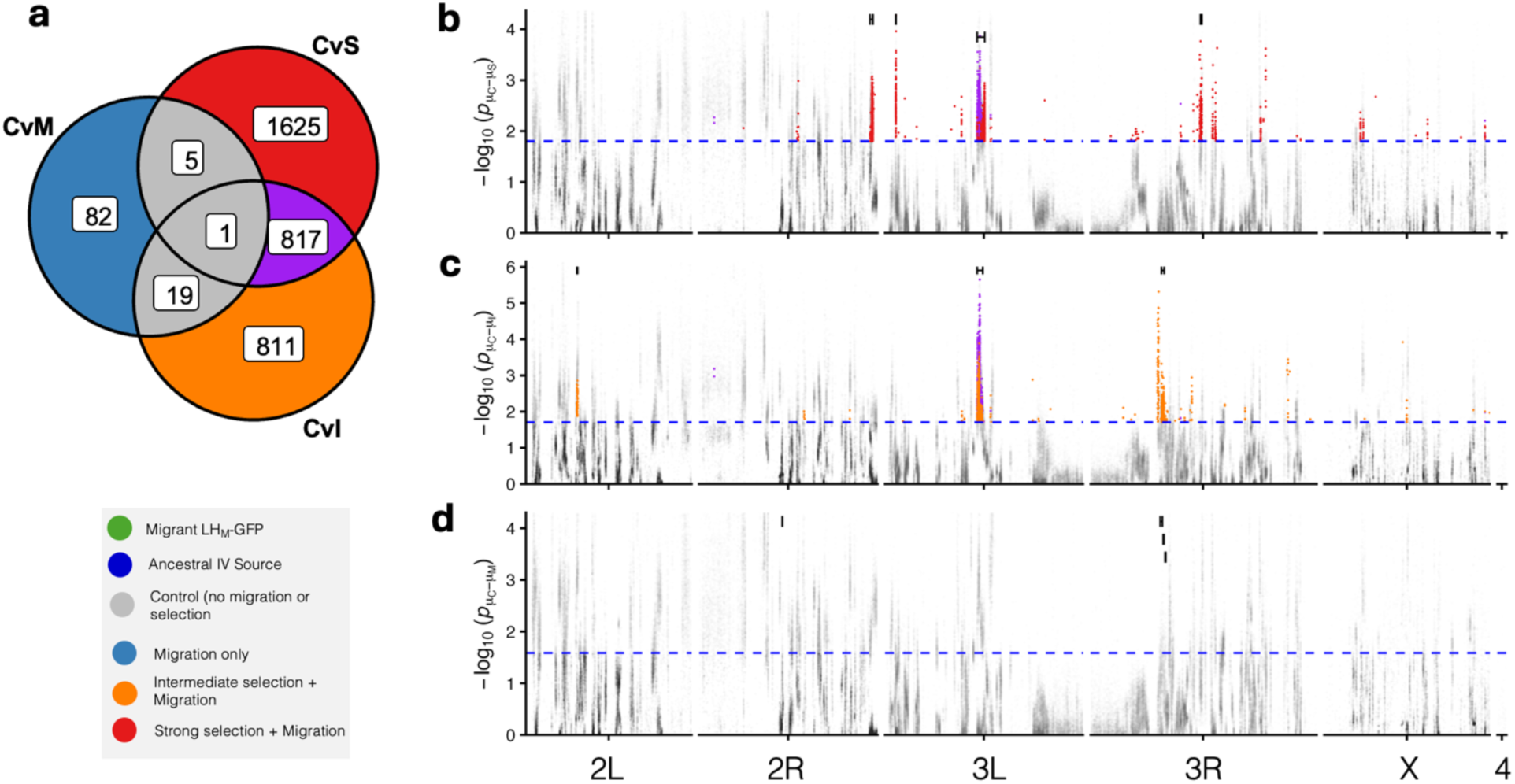
SNPs showing a signature of divergent selection across all major *D. melanogaster* chromosomes. (a) Venn diagram showing the overlap in loci displaying a pattern of divergent selection of the I, S, and M contarasts across all chromosomes. (b-d) Manhattan plots displaying the most significant divergent selection SNPs in the (b) C-S, (c) C-I, and (d) C-M tests. Dotted blue line denotes FDR threshold of 0.05. Brackets bound highly correlated SNPs assigned to independently evolving clusters using a predefined distance and *rw* threshold (Methods). Red points indicate S-exclusive SNPs, orange points indicate I-exclusive SNPs, blue points indicate M-exclusive SNPs, and purple points indicate SNPs shared between I and S.

**Extended Data Figure 10:**
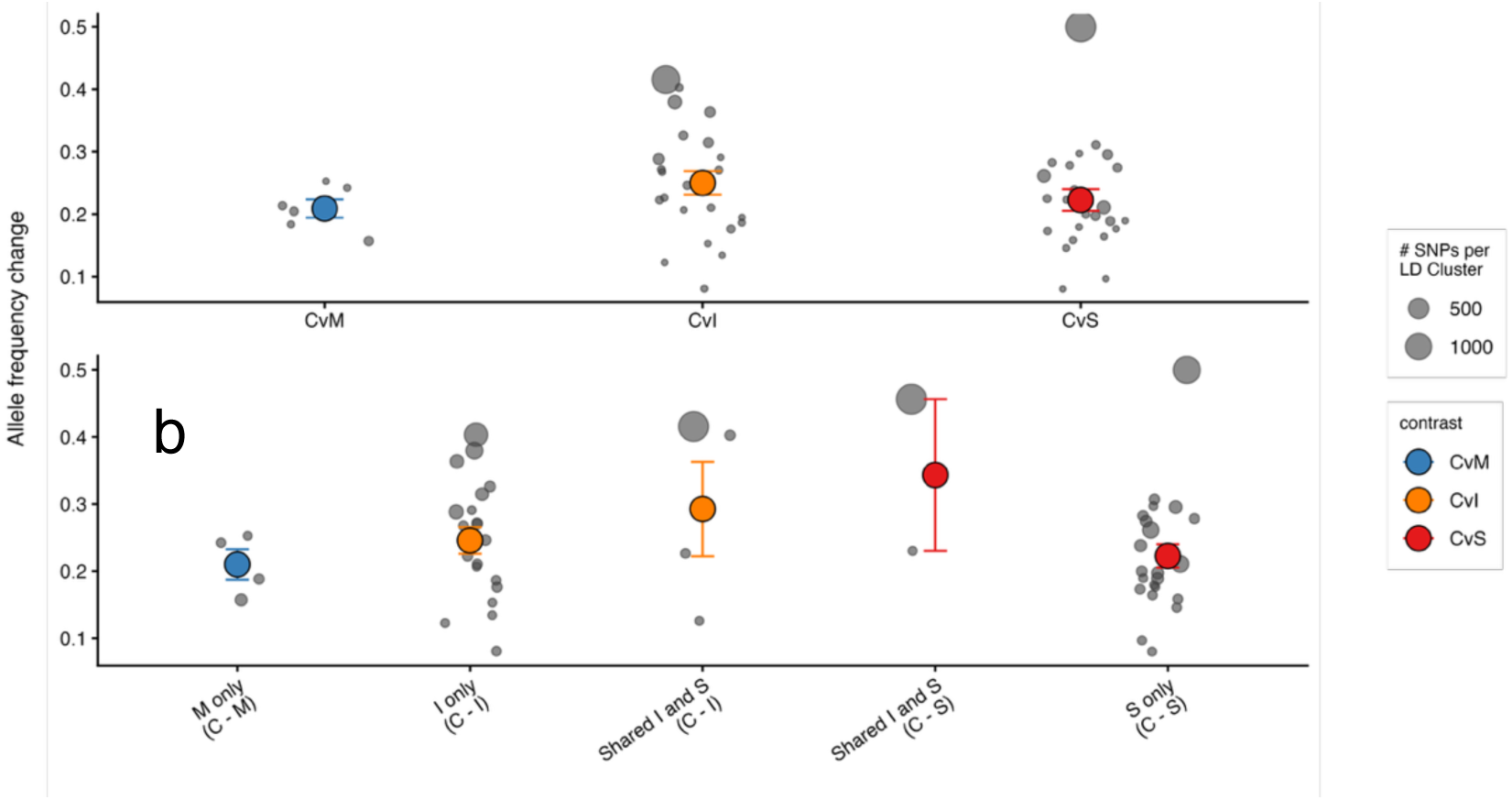
(a) Average allele frequency change from C of significant divergent selection candidate SNPs on chromosome 3 grouped by contrast. (b) Average allele frequency change from C emphasizing SNPs with shared patterns in C-I and C-S. Each point represents an independent SNP cluster. Point size of grey points represents number of SNPs assigned to that cluster. Clusters with only a single SNP were excluded from the analysis. Colored points represent mean estimate of each peak SNP (SNP with the largest change from C) per cluster assigned to each contrast +/− 1 S.E.

